# Structural Knowledge Transfer of Panoptic Kidney Segmentation to Other Stains, Organs, and Species

**DOI:** 10.1101/2021.10.21.465370

**Authors:** Brandon Ginley, Kuang-Yu Jen, Pinaki Sarder

## Abstract

**Background:** Panoptic segmentation networks are a newer class of image segmentation algorithms that are constrained to understand the difference between instance-type objects (objects that are discrete countable entities, such as renal tubules) and group-type objects (uncountable, amorphous regions of texture such as renal interstitium). This class of deep networks has unique advantages for biological datasets, particularly in computational pathology.

**Methods:** We collected 126 periodic acid Schiff whole slide images of native diabetic nephropathy, lupus nephritis, and transplant surveillance kidney biopsies, and fully annotated them for the following micro-compartments: interstitium, glomeruli, globally sclerotic glomeruli, tubules, and arterial tree (arteries/arterioles). Using this data, we trained a panoptic feature pyramid network. We compared performance of the network against a renal pathologist’s annotations, and the method’s transferability to other computational pathology domain tasks was investigated.

**Results:** The panoptic feature pyramid networks showed high performance as compared to renal pathologist for all of the annotated classes in a testing set of transplant kidney biopsies. The network was not only able to generalize its object understanding across different stains and species of kidney data, but also across several organ types.

**Conclusions:** Panoptic networks have unique advantages for computational pathology; namely, these networks internally model structural morphology, which aids bootstrapping of annotations for new computational pathology tasks.

## Introduction

A plethora of studies have shown the undeniable utility of deep neural networks for the detection of histopathological structures in digital pathology datasets.^1-10^ However, many earlier works commonly employed pure semantic segmentation, or pure instance segmentation, missing out on the best qualities of both. Semantic segmentation is the task of assigning a classification label to every pixel in the image, which is flawed in that it cannot distinctly recognize two same-class entities that are abutting or overlapped in an image. Instance segmentation is the task of distinctly recognizing abutting/overlapping objects as unique entities, but typically does not model multiple classes.

Panoptic segmentation was designed to unify the tasks of instance and semantic segmentation, and allows the detection of multiple classes and their boundaries in a single model.^11-13^ Panoptic segmentation tasks require target regions of interest to be defined either as group-type objects (colloquially termed “stuff”, uncountable amorphic regions of texture like water or sky) or as instance-type objects (colloquially termed “things”, countable objects with distinct boundaries like people or cars). This strategy is particularly advantageous for biological studies, which often desire to measure both multiple instance classes (e.g., nuclei, glomeruli, and tubules) and multiple semantic classes (e.g., cortex, medulla, and interstitial fibrosis and tubular atrophy (IFTA)). Moreover, the combined modeling of the panoptic network eliminates the need to explicitly annotate the divider between classes, as was done in prior studies.^2^

To investigate the performance of panoptic segmentation for digital pathology segmentation tasks, we trained a panoptic feature pyramid network (pFPN) on a large dataset (*n* = 126) of periodic acid-Schiff (PAS) stained human kidney biopsies, fully annotated for interstitium (group type object), glomeruli, sclerotic glomeruli, tubules, and the arterial tree (instance type objects). We next evaluated the performance of this trained model in a holdout set of ten transplant kidney biopsies, one-half of which were histologically-normal (as per renal pathologist visual assessment), and the remaining half were diseased. The network achieved excellent performance on each tested image class, with the lowest scoring class being the arterial tree achieving a dice coefficient of 0.93.

Upon achieving an ideal computational performance for human PAS kidney tissue sections, we next studied the ability of the network to understand objects in new stains, species, and organs, *without any retraining*. We found that the pFPN model generalizes to segment the aforementioned five kidney classes in hematoxylin & eosin (H&E) stained human kidney biopsies and PAS stained murine kidney biopsies. Moreover, the network was capable of identifying bile ducts in both H&E and trichrome stained human liver, and glands or other bounded cellular units in H&E stained stomach, small intestine, pancreas, appendix, large intestine, and rectum. The network segmentation of single objects from non-kidney organs was found to be precise, suggesting they could be reused for the purpose of bootstrapping new digital pathology models.

Our contributions in this work are: (i) Densely annotating a large set of kidney biopsies for training of segmentation networks, (ii) optimizing convolutional network input and hyperparameters for kidney segmentation, (iii) training and evaluating a panoptic segmentation network for kidney histopathology analysis, and (iv) evaluating transferability of convolutional predictions to diverse tissues.

## Methods

### Image data

Whole slide images (WSIs) used in this study were sectioned at 2-3 µm, stained with various stains, including periodic acid-Schiff (PAS), Hematoxylin and Eosin (H&E), Jones silver, or trichrome, and scanned either at 40x (kidney, liver samples, 0.25µm/pixel) or 20x (all other organs, 0.5µm/pixel) by a whole slide brightfield microscopy image scanner (Aperio, Leica).

### Training

126 WSIs of PAS stained kidney biopsies were fully annotated for interstitium, glomeruli, globally sclerotic glomeruli, tubules, and the arterial tree (arterioles and arteries), and stored as Aperio^®^ ImageScope generated XML files. Each WSI in the training set was gridded at high resolution into 1200 × 1200 non-overlapping blocks.

Tiles containing less than 5% tissue were excluded. To identify these tiles, first, a low-resolution thumbnail of the WSI image was extracted, and transformed with an HSV (hue, saturation, value) color space transformation. Pixels constituting tissue region were defined as those pixels with saturation value greater than 0.05, which yields a low-resolution binary mask of the usable regions. Any tiles with the sum of usable regions less than <5% of the total tile area were excluded from processing. Corresponding image masks for network training were created by parsing the XML file for the associated coordinates of contours within the tile. A total of 113,841 unique non-overlapping tiles were extracted for training.

Network training was orchestrated using the Dectron2 library for PyTorch,^14^ which implements convenient functions for training and evaluating a pFPN architecture. Network weights were initialized to a model pretrained on the COCO dataset^15^ available in the Detectron2 library, which had ResNet-50 backbone and was originally trained with a 3X learning rate schedule. For re-training, a similar training schedule was followed as was laid out in the original implementation^12^ for training on the COCO dataset. Starting from the COCO pretrained model, we trained the pFPN for a total of 296K steps, with a step learning rate policy, starting at 0.02, and dropping by one tenth upon reaching 182K and 250K steps, respectively. The batch size was set to four, in order to train on a single GeForce RTX 2080 TI (11GB of memory). Glomeruli (sclerotic and not), tubules, and the arterial tree were specified as instance-type segmentation objects (“things”), and the interstitium and background were specified as group-type segmentation objects (“stuff”). The batch size per ROI head was set to 128 to reduce memory. The default behavior of Dectron2 to filter empty annotations was turned off, as many unlabeled useful training patches exist in our dataset. Any other training parameters were left as the default configuration specified by the pFPN implemented in Dectron2.

### Inference

Testing was performed on 10 WSIs. For each WSI, the entire biopsy region was tiled into 3000 × 3000 blocks, each of which was input to the network for prediction. The resultant outputs were re-stitched into a WSI prediction mask using the original chopping coordinates. One typical challenge in this process is the presence of network output edge artifacts induced by the tiling process.^16-18^ Namely, chopping the WSI into a grid of patches results in many image patches with structures that are not fully contained within the patch. This becomes an issue due to the mathematical formulation of convolutional operations, which rely on pixel neighborhood information to extract meaningful features, and pixels on the image edge do not have full neighborhood information. We found that selecting our patch size to be 3000 × 3000 reduces the total amount of tiling needed, thereby reducing the tiling artifact. Further, our testing patch-grids overlap 400 pixels between adjacent tiles. When stitching the prediction masks into a larger WSI prediction mask, the trailing and leading 200 pixels of adjacent prediction patches are clipped, resulting in a final WSI prediction mask free of any tiling artifacts. WSI prediction mask objects are converted to a series of contours and stored as an XML file.

### Performance analysis

Ten WSIs from transplant surveillance kidney biopsies were used for testing, five from patients which had serum creatinine at time of biopsy >2, and five from patients with “histologically normal” biopsy findings as per the renal pathologist. This strategy was followed to test the performance both in disease and in healthy data. Image data for ground-truth annotation was selected by randomly sampling three tiles sent to the network for prediction from each of the 10 testing WSIs, resulting in a total of 30 large image patches. Testing images containing majority medulla or slide background were re-sampled until the patch contained sufficient cortical region for analysis. These cropped sub-regions were converted to tiff images and provided to the pathologist who created the ground truth annotation for performance evaluation using Aperio^®^ ImageScope. From each pair of image predictions and annotations, sensitivity, specificity, precision, negative predictive value, Matthew’s correlation coefficient, and the dice coefficient were calculated, using a *one-versus-all* approach for glomeruli, sclerotic glomeruli, tubules, and the arterial tree. Specifically, for each class, we first pool all true and false, positive and negative pixels from all 30 test images, then calculate the performance metrics one time for the entire set of testing pixels. Performance was not assessed for interstitium class as it is a catch-all category for remaining structures which are not directly used in clinical assessments, and simply sets a base outline for future sub-compartmentalization of this region.

## Results

Figure 1 shows the performance of the pFPN to segment renal structures in a PAS stained kidney biopsy from the test set. Fig. 1A shows the network prediction thumbnail, where the prediction boundaries are so dense they are barely discernable. Fig. 1B shows a zoomed magnification of the network outputs, showing the network to have excellent detection of all target kidney classes (globally sclerotic glomeruli not seen in magnified view, although the sole sclerotic glomerulus in this biopsy can be seen south of the left-most inset). The interstitium class correctly avoids perivascular stroma, which was our intention upon creating the training data. Fig. 1C shows another magnified region which, from low resolution, appears to be a large artifactual false positive. However, upon magnifying, it becomes clear it is a partially segmented large caliber artery. This is an intuitive result as arteries this size are scarce in kidney biopsy and thus scarce in the training set. Additionally, though our training set was constructed of only biopsies with >90% cortex, the network can be observed to have an excellent understanding of arterioles and tubules in the medulla.

**Figure 1.**
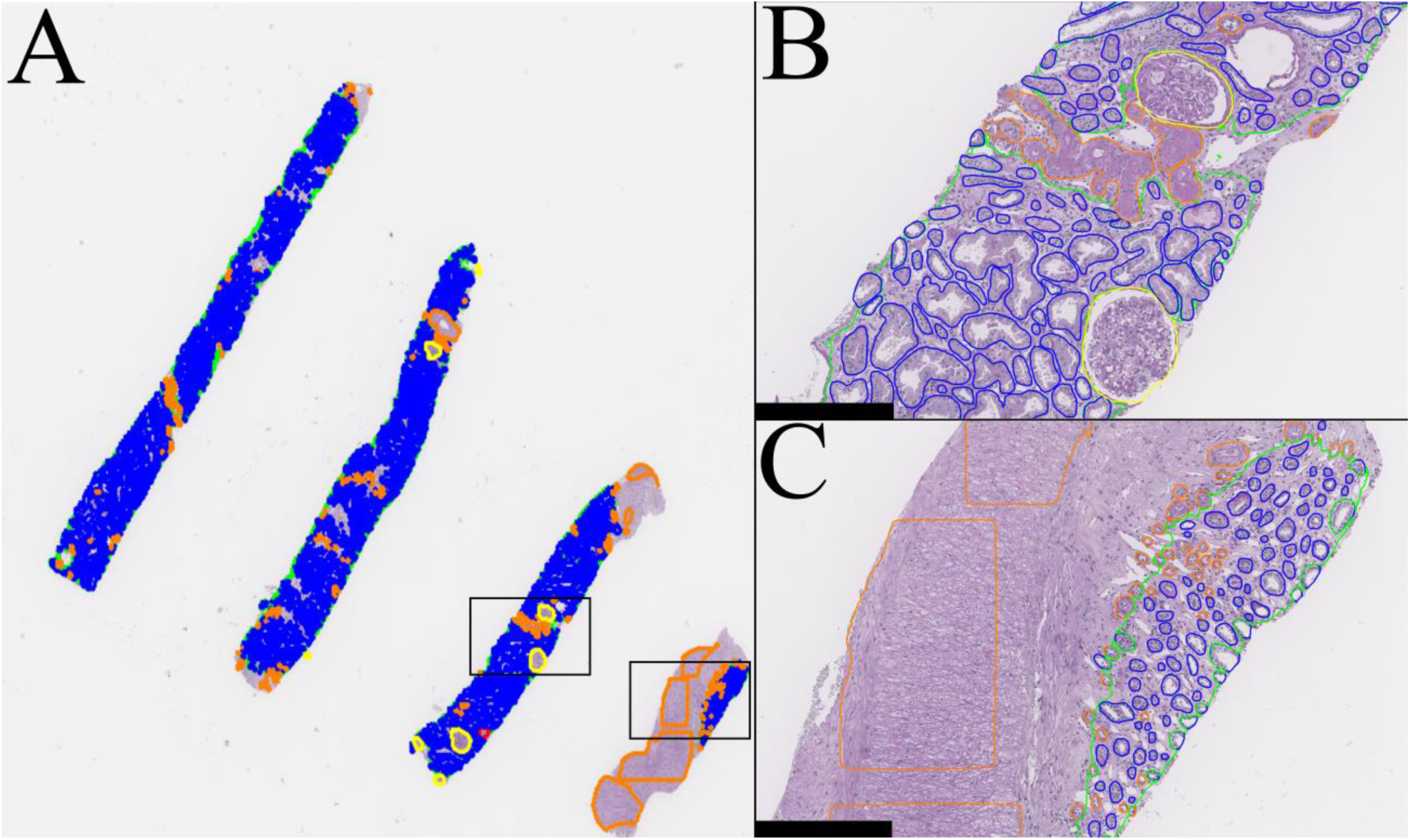
Output whole biopsy segmentations for a human, PAS-stained transplant kidney biopsy from the testing set. A) Thumbnail image of segmentation boundaries for the entire biopsy image (not to scale). B) 10x magnification zoom on the left inset of panel A. C) 10x magnification zoom on the right inset of panel A. Prediction colors: green-interstitium, yellow-glomeruli, red-globally sclerotic glomeruli, blue-tubules, orange-arterial tree.

Table 1 shows the network’s performance compared pixel-wise against renal pathologist ground truth. Glomerular classes showed extremely high performance, and we did not observe any incorrectly predicted glomerular boundaries in the testing set. This result is reinforced by the network’s ability to detect glomeruli effectively in other stains and species, see Figure 2. The next best performing class was tubules, again very intuitive due to their relatively simple morphological structure and ample presence in the training set. The arterial tree class performed worst (precision 0.88), due to its inability to effectively identify small diameter arterioles, or those cut in tangential patterns. This result may be because smaller/tangentially cut arterioles are challenging to identify even by manual means, and can be relatively rare compared to other image classes. However, the performance values are still high, due to its accurate segmentation of larger caliber arterioles and arteries with normal cross-sectional profiles, which are most commonly assessed for interpretation of renal vascular disease.

**Table 1.** Performance metrics compared against renal pathologist.

|  | Sensitivity | Specificity | Precision | NPV | MCC | Dice |
| --- | --- | --- | --- | --- | --- | --- |
| Glomeruli | 0.99 | 1 | 0.99 | 1 | 0.99 | 0.99 |
| Sclerotic glomeruli | 1 | 1 | 1 | 1 | 1 | 1 |
| Tubules | 1 | 0.95 | 0.91 | 1 | 0.93 | 0.95 |
| Arterial tree | 0.99 | 1 | 0.88 | 1 | 0.93 | 0.93 |

NPV: Negative predictive value, MCC: Matthew’s correlation coefficient

**Figure 2.**
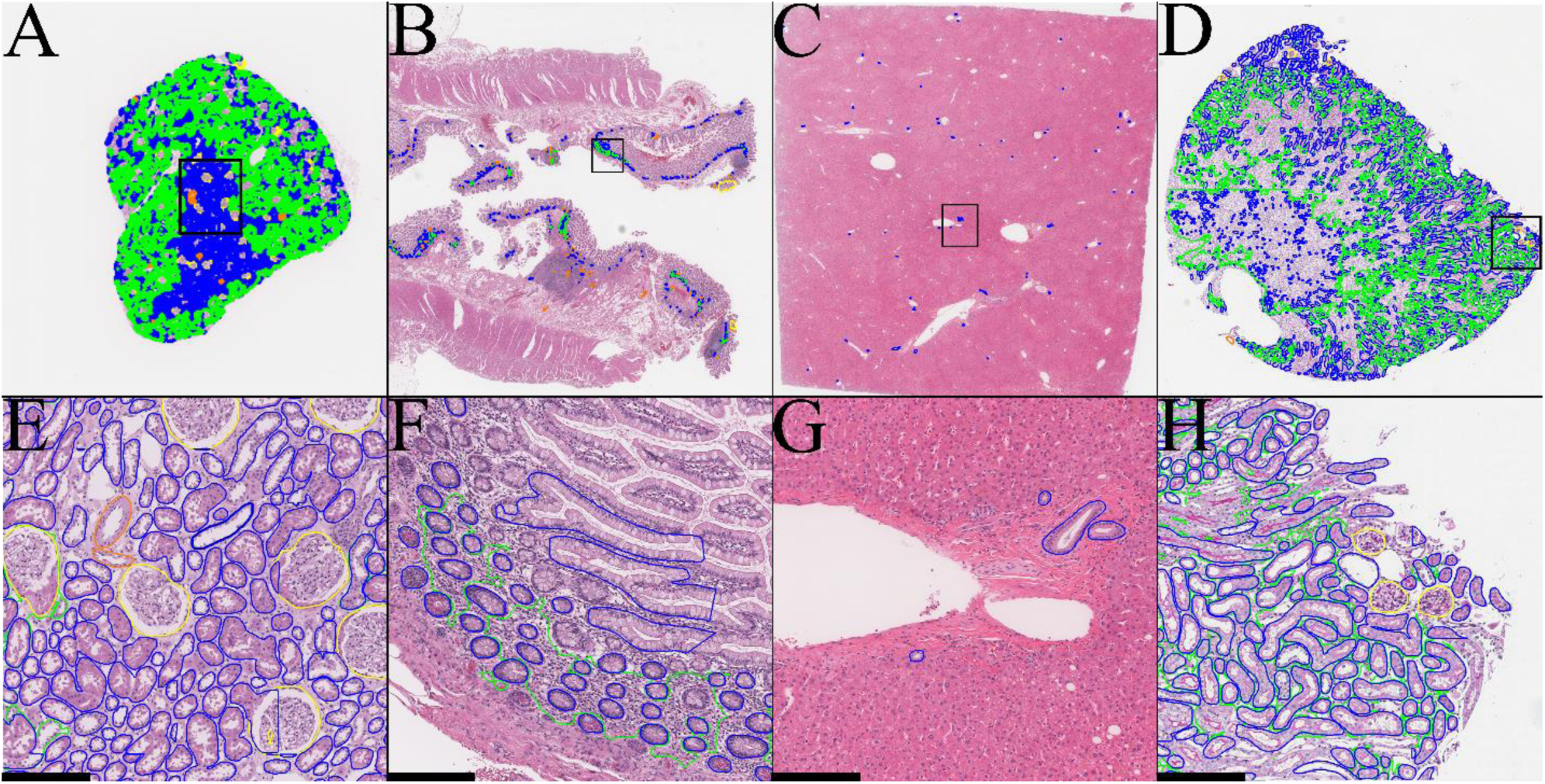
Untrained transfer of structure knowledge across stain, species, and organ. Green, blue, yellow, and orange correspond to network predictions of interstitium, tubules, glomeruli, and the arterial tree, respectively. Figs. A-D) Show respective thumbnail images of entire slide predictions (images not to scale) for H&E kidney, H&E stomach, H&E liver, and PAS murine kidney, respectively. Figs. E-H) Show zoomed insets (10x) of the respective black outlined regions in A-D (scale bar 200 µm).

Figure 2 shows the ability of the network to generalize histological structures beyond stain, species, and organ. The network transferred its structural understanding optimally to human H&E kidney tissues, capable of segmenting all five classes, albeit inaccurate in some regions (Fig. 2A/2E). The network was also able to identify glands within an H&E stained small intestine biopsy (Fig. 2B/2F), and bile ducts within an H&E stained liver biopsy (Fig. 2C/2G). In all cases, the network demonstrates a clear generalized understanding of tubular cellular organization in other human organs. The network was also capable of segmenting all five classes in PAS stained murine kidney tissues (Fig. 2D/2H), though a much larger number of missed tubules was observed compared to the human tissue. Surprisingly, murine glomeruli were segmented perfectly. Similar predictions were observed for Jones stained human kidney, trichrome stained human liver, as well as H&E biopsies from human stomach, rectum, pancreas, appendix, and large and small intestines. These predictions are extremely useful in reducing annotation burden for computational pathology tasks.

An example routine to bootstrap a new organ segmentation model based on our results would be as follows: first, select several ROIs, such as those identified in Fig. 2E-H, such that approximately 50% or more of the desired structures are correctly predicted. This percentage can be scaled up or down depending on available annotation effort. On one hand, having more correct predictions means less annotation effort, but on the other hand, correcting more of the improper segmentations will lead to better network output (termed active learning^19-21^). Then, the corrected ROIs should be mixed with the already established training set, reassigning classes of different organ systems into a discrete ontology,^22^ where needed. Additionally, any ROIs selected for retraining should ideally be captured from different WSI cases to maximize training diversity, and therefore benefit, from the retraining process.

## Conclusions

Despite only being trained on PAS stained human kidney tissue, our pFPN model was capable of detecting target classes in other stain types, such as H&E, and kidney from other species, such as mice. Moreover, without retraining, the pFPN was capable of generalizing its structural understanding of tubules in kidney to bile ducts of H&E liver and glands of H&E small intestine. This structural generalization without training is extremely valuable for computational pathology tasks, which often require extreme annotation effort at their genesis. Further, panoptic networks offer a unique opportunity for biomedical researchers to inject ontological schemas into the network modeling process.

## Author contributions

BG conceptualized and implemented the study design, generated multi-compartment training annotations verified by KYJ prior to network training, performed all study analysis, and wrote the manuscript. KYJ provided slide data, training annotations, performance evaluation annotations, and clinical feedback on the overall work and manuscript. PS coordinated the study organization and resources to perform the study, and gave feedback on the study design and manuscript preparation.

## Acknowledgments

This project was supported by NIH-NIDDK grant R01 DK114485 (PS), NIH-OD grant R01 DK114485 03S1 (PS), a glue grant (PS) of the NIH-NIDDK Kidney Precision Medicine Project grant U2C DK114886 (Contact: Dr. Jonathan Himmelfarb), and NIH-OD grant U54 HL145608 (PS).

## Notes

### Competing Interest Statement

The authors have declared no competing interest.

